# Tuning of B_12_ photochemistry in the CarH photoreceptor to avoid radical photoproduct

**DOI:** 10.1101/2023.08.11.552799

**Authors:** Ines S. Camacho, Emma Wall, Igor V. Sazanovich, Emma Gozzard, Mike Towrie, Neil T. Hunt, Sam Hay, Alex R. Jones

## Abstract

Time-resolved infrared spectroscopy reveals the flow of electron density through coenzyme B_12_ in the light-activated, bacterial transcriptional regulator, CarH. The protein stabilises a series of charge transfer states that result in a photoresponse that avoids reactive, and potentially damaging, radical photoproducts.

---

The bacterial transcriptional regulator, CarH, employs the photochemistry of coenzyme B_12_ (5’-deoxyadenosylcobalamin, AdoCbl, Fig. S1) to trigger the biosynthesis of carotenoids in response to photooxidative stress.^1, 2^ Binding of AdoCbl to CarH from *Thermus thermophilus* for example, drives the formation of homotetramers,^3^ which in turn bind to and block operator DNA.^4^ Photodissociation of the upper axial 5’-deoxyadenosyl (Ado) ligand^5^ by absorption of wavelengths below 600 nm results in DNA release following disassembly of the tetramer^4^ into CarH-monomer / Cbl adducts.^6^ The fact that the CarH photoresponse enables both gene activation and changes to protein oligomerisation state has been exploited in various optogenetic^7, 8^ and smart material^9^ applications.

Typically, AdoCbl participates in enzyme mechanism^10^ and photochemistry^2^ that proceed *via* cob(II)alamin / Ado radical pair intermediates. By contrast, CarH appears to tune the photochemistry of AdoCbl away from radicals,^5^ which is perhaps appropriate for a protein that responds to photooxidative stress and is bound to DNA. UV-visible transient absorption (TA) spectroscopy has revealed that the productive photochemical channel proceeds *via* cob(III)alamin intermediates,^5, 11^ which are preceded by a metal-to-ligand charge transfer (MLCT) state similar to that observed following photoexcitation of methylcobalamin (MeCbl).^12^ The alkyl photoproduct from this process is 4′,5′-anhydroadenosine,^13^ instead of the highly reactive primary carbon Ado radical.

The details of how CarH tunes the photochemistry of AdoCbl are currently unknown but are vital to our understanding of its natural function and for the optimisation of genetically-encoded tools based on B_12_-dependent photoreceptors. Time-resolved infrared (TRIR) spectroscopy is a powerful method to relate mid-IR signals from the protein scaffold to the photophysical and photochemical dynamics of bound chromophores. Moreover, it has been used successfully to investigate the role of local protein dynamics in AdoCbl-dependent enzymes.^14-16^ From TRIR data acquired up to microseconds with femtosecond resolution, here we show that CarH stabilises the charge transfer (CT) states of AdoCbl that steer its photochemistry away from radical photoproducts.

## CarH is a challenging system for study using TRIR

The CarH tetramer is sufficiently soluble at concentrations required to generate good quality TRIR signals (∼ 250 μM monomer equivalent, Fig. 1). Solubility at these concentrations is reduced in its monomeric state, however, which leads to precipitation upon photoexcitation. This presents a challenge because protein sample volumes are usually limited and are therefore typically cycled through the optical path during TRIR data acquisition to enable data averaging for improved signal-to-noise ratio. In the case of CarH, sample cycling would result in scattering of the probe light by precipitated photoproduct, which would obscure the signal.

**Fig. 1.**
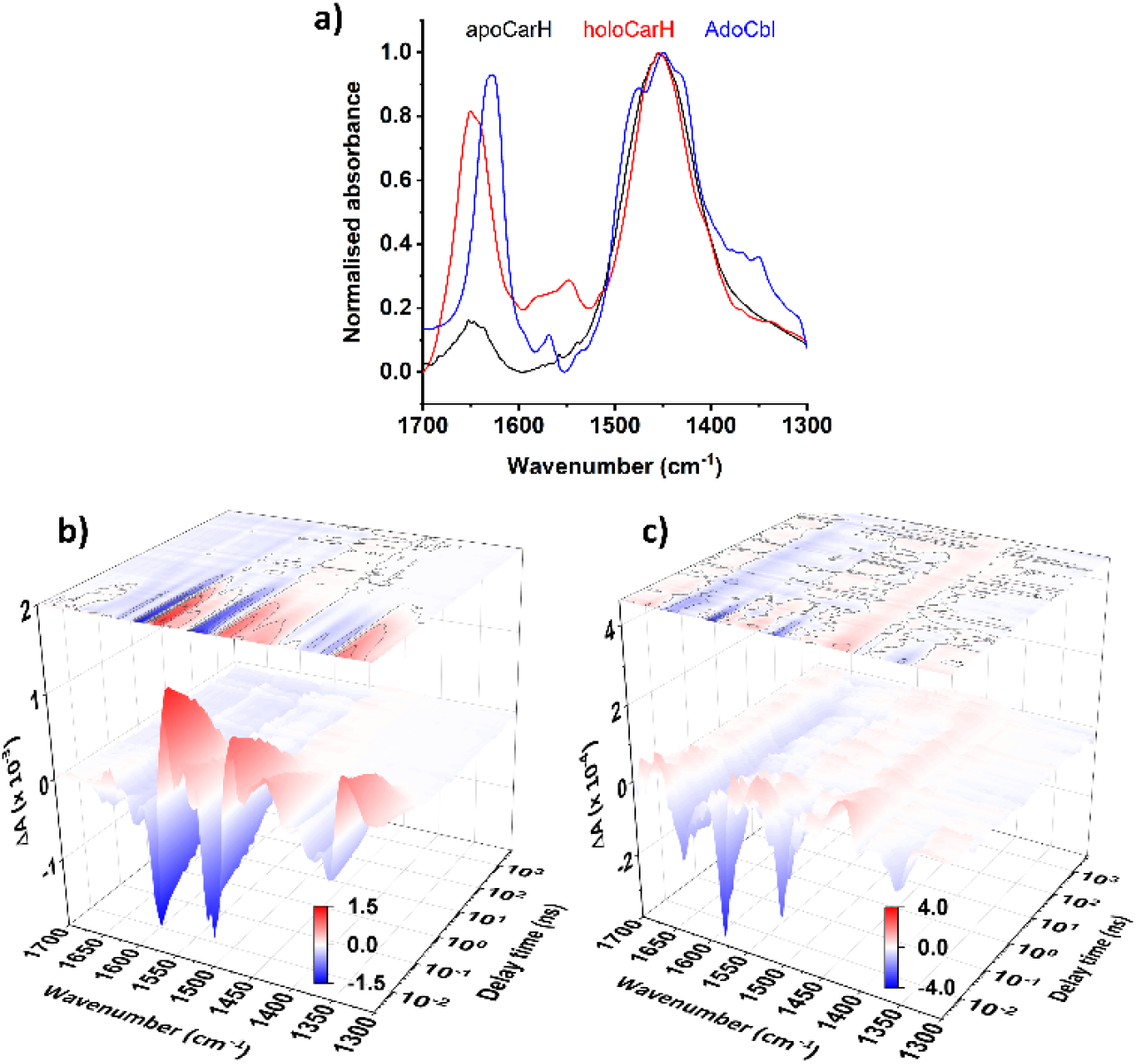
**a)** Normalised FTIR absorption spectra of AdoCbl and CarH (apo- and holo-protein). TRIR data matrices between 2 ps – 5 μs following the photoexcitation at 525 nm of ∼3 mM free AdoCbl **(b)** and ∼250 μM holoCarH **(c)**. False colour scales in (b) and (c) are the same as the respective y-axis (10^−^3 & 10^−4^, respectively).

To prevent degradation of data quality by photoinduced precipitation, and confounding signals from any soluble photoproduct,^5^ we developed a bespoke sample flow system (described in the Supplementary Methods and elsewhere^17^). Briefly, the sample was flowed one-way at an optimised rate through an optical cell during data acquisition using a syringe pump (Fig. S2). The sample was then collected, and any precipitated photoproduct removed *via* centrifugation. The supernatant was reloaded into the syringe and flowed again for further data acquisition. Because only a small proportion was photoconverted in each pass, the sample losses were minimal, and it was possible to collect at least two datasets from each sample (Fig. S3). We thus collected high quality TRIR spectral data at probe wavenumbers between 1700 – 1300 cm^−1^ for delay times ranging from 2 ps to 5 μs (Figs. 1&S4).

### CarH tunes the electronic and vibrational structure of AdoCbl

The ground state Fourier transform IR (FTIR) absorption spectra of AdoCbl, the CarH apoprotein (apoCarH), and the CarH holoprotein (holoCarH) are broadly similar with slight variations (Fig. 1a). Each has a broad feature in the amide II’ region (∼ 1450 cm^−1^). The minor peak at 1569 cm^−1^ in the spectrum of free AdoCbl is absent for apoCarH and shifted to 1548 cm^−1^ when AdoCbl is bound to CarH. This signal corresponds to the corrin breathing mode and is very sensitive to the electronic structure of the axial ligation.^14, 18^ The observed shift will in part be owing to the displacement of the lower axial 5’,6’-dimethyl-benzimidazole ligand in AdoCbl by His177 when bound to CarH. It could also indicate changes to the electronic structure of the upper axial Ado, which we explore below. There are peaks in the amide I region at 1630 cm^−1^ and 1646 cm^−1^ in the spectra of free AdoCbl and holoCarH, respectively. The band in AdoCbl may be expected to shift to lower frequencies when bound to a protein, because H bonds appear to stabilise the charged resonance form of the corrin propionamides.^18^ Inspection of the CarH crystal structure, however, suggests that only two of the six corrin propionamides are likely to form H bonds with the protein in CarH (Fig. S5a).^6^ The shifting to higher frequency in holoCarH is therefore likely owing to the amide I signal from the protein (which peaks at 1647 cm^−1^ for apoCarH, Fig. S5b).

### Binding of AdoCbl to CarH has a significant impact on the TRIR signals compared to AdoCbl free in solution (Figs. 1b&c)

Our previous studies characterised the transient mid-IR spectral fingerprints for a range of B_12_ derivatives.^14^ Briefly, following photoexcitation with green light (525 nm), free AdoCbl produces difference signals across the 1700 – 1300 cm^−1^ within 2 ps. Most of the amplitude decays over the course of a few ns to leave residual signals corresponding to the longer-lived, solvent-separated radical pairs, consistent with published UV-visible TA data.^19^ The difference spectra change little during this time apart from some minor peak shifts. Density functional theory (DFT) calculations reveal the bands that shift (*e*.*g*., the transient at ∼ 1550 cm^−1^) correspond to coupled motions through the corrin macrocycle, and the kinetics indicate the changes correlate with reduced steric interaction with the bulky Ado radical during diffusive separation of the radical pair.^14^ The free AdoCbl TRIR data acquired here are consistent with these observations (Figs. 1b&S4). The small size of the differences suggests that, vibrationally, the initial excited states of free AdoCbl closely resemble the radical pair relative to the ground state. This conclusion is supported by DFT calculations from our previous TRIR study of B_12_ cofactors.^14^

In CarH (Fig. 1c), apart from the early bleaches (negative signals) that represent the depopulation of the ground state, the signals and their dynamics are different from free AdoCbl, with no evidence of radical photoproducts. The transients (positive signals) deviate from those observed for free AdoCbl and are supressed in amplitude relative to the bleaches. The decay of the initial signals is also much more rapid in CarH (within a few hundred ps) leaving a spectrum with major bands that peak at 1633 and 1423 cm^−1^, which persists throughout the 5 μs acquisition window. The nature of the differences in photoresponse between CarH and free AdoCbl are revealed in the evolution associated spectra (EAS, Fig. 2) from global kinetic analysis. The spectra and kinetics appear to be independent of excitation wavelength for both AdoCbl and CarH (Figs. S6&S7). The one exception is, when photoexcited at 380 nm (Fig. S7), the quantum yield of the CarH photoresponse is much reduced compared to photoexcitation at 525 nm (Fig. 1c), as reported elsewhere.^11^

**Fig. 2.**
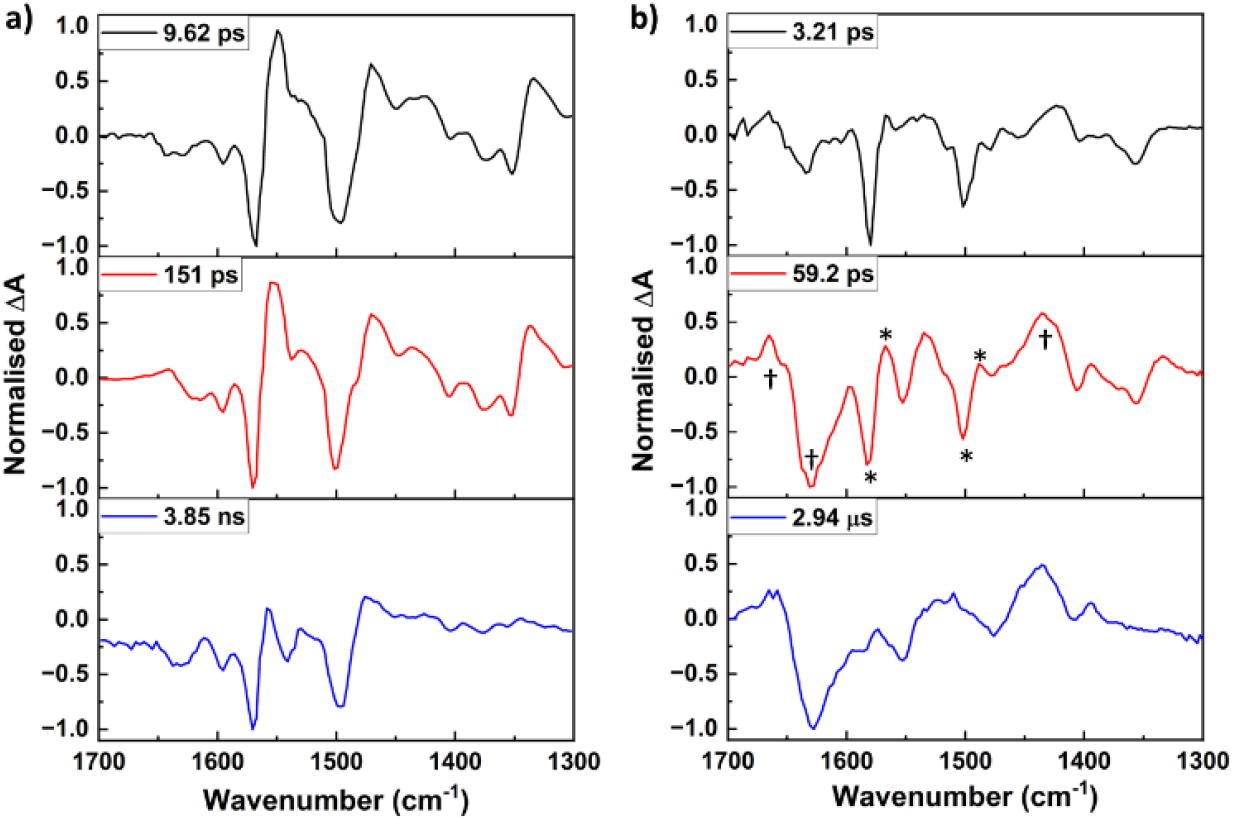
Evolution associated spectra (EAS) and respective lifetimes from the global analysis of TRIR data from free ∼3 mM AdoCbl **(a)** and ∼250 μM CarH holoprotein **(b)**. Y-axes are normalised between −1 and 1; notable signals are highlighted with * and † in (b) and discussed in the text. See SI for uncertainty / sensitivity analysis.

### The initial excited state of CarH is predominantly a ligand-to-metal charge transfer state (LMCT)

The EAS for the initial excited state from the UV-visible^19, 20^ and TRIR (Fig. 2a) data from free AdoCbl have been assigned to a MLCT state, where electron density is transferred from the Co-axial ligation to the corrin π* orbitals.^21^ The EAS of the equivalent state in CarH from the TRIR (Fig. 2b) and UV-visible^5^ data are quite different and closely resemble those of the CNCbl excited state (Fig. S8).^14, 22^ Experiment^22^ and theory^23^ have assigned the lowest lying excited state of CNCbl as a π → 3dz^2^ LMCT (corrin to Co) state, where both axial bonds to the cobalt are weakened and lengthened, with contributions from a ligand-to-ligand (corrin to CN) CT state.^24^ The axial elongation that accompanies the LMCT state in CNCbl means the barrier to internal conversion back to the ground state is low,^23^ hence CNCbl is photostable and shows a single decay component in UV-visible TA^22^ and the TRIR data (Fig. S8).^14^ By contrast, this excited state in CarH converts to subsequent intermediates (Fig. 2b). Recent time-resolved X-ray absorption near-edge structure (XANES) data suggest CarH limits the axial elongation,^11^ which presumably increases the barrier to internal conversion relative to CNCbl. In both CNCbl and CarH, stabilisation of this LMCT state appears to disfavour formation of radical photoproducts.

### TRIR signals consistent with radical pair intermediates in CarH are short-lived

The EAS from global analysis of the CarH TRIR data over 5 μs reveal two further components following the initial excited state. Both have shared features with additional signals only in the second EAS (marked † and *, respectively, Fig. 2b). The latter signals strongly resemble those from radicals generated following photolysis in free AdoCbl (second EAS, Fig. 2a), exemplified by the bleaches at ∼ 1500 and 1570 cm^−1^. Consistent with previous UV-visible TA data from CarH,^5^ these radical signals decay with a lifetime of ∼ 60 ps. It is therefore likely that the second EAS in Fig. 2b is a convolution of two species (Fig. S9a), with the radical signals rapidly decaying to leave the intermediate represented by the third EAS. The amplitude of the CarH signal is low, particularly at longer delay times, and an improved S/N might be required to separate the convoluted signals. Despite this, our sensitivity analysis (see Supplementary Information) suggests the kinetics are well described. In [5], we proposed a branching mechanism, where the radical species are part of a low-population, legacy channel, which would be consistent with convolution of signals. We explored both sequential and branching models in our analysis, however, and they made little difference to the outcome (Figs. S9b&c). We touch on this further below. The sub-ns radical pair recombination kinetics observed here and in [5] for CarH is much faster than for AdoCbl free in solution and is presumably owing to the protein serving as a cage that to some extent limits radical diffusion.

### The CarH TRIR data contain long-lived signals from the protein

The third EAS from global analysis of the CarH TRIR data contains several broad features, which decay with a 2.9 μs lifetime: bleaches peaking at 1625, 1553 and 1480 cm^−1^, and transients peaking at 1662, 1516 and 1435 cm^−1^. In most respects, this is unlike anything from similar measurements with B_12_ derivatives.^14^ The only similarities are the transient (1662 cm^−1^) and broad bleach (1625 cm^−1^) in the protein amide I region, which resemble mid-IR signals from protein motions coupled to catalysis by the AdoCbl-dependent enzyme, ethanolamine ammonia lyase.^15^ Moreover, we noted above that the amide I band in the holoCarH FTIR spectrum (Fig. 2a) is likely dominated by signal from the protein. It appears likely, therefore, that these long-lived spectral components from the CarH TRIR data represent changes to the protein scaffold following photoexcitation of AdoCbl. This is also supported by the recovery of the ground state bleaches from AdoCbl.

### Are these long-lived signals relevant to the productive CarH photoresponse

or do they represent heat dissipation through the system^25^ following photoexcitation? To investigate, we conducted variable temperature FTIR measurements for AdoCbl, holoCarH, and apoCarH. Spectra were acquired between 1700 – 1300 cm^−1^ for each sample every 2 degrees between 22-40 °C, with a corresponding spectrum acquired at 20 °C subtracted from each to assess temperature-induced changes (Fig. S10). These were then corrected for signal changes from the solvent (Figs. S10&S11). The data from the largest temperature difference for holoCarH were compared to the 3^rd^ EAS from the global analysis of the CarH TRIR data (Fig. S11) and difference spectra at selected time delays from the raw data (Fig. S12). It is clear from these comparisons that heat dissipation can account for only some of the signals represented by the third EAS. Of note are the transient and bleach identified above in the amide I region. No equivalent signals to these are evident in any of the difference FTIR spectra following heating (Fig. S11) and are therefore unique to the holoCarH TRIR data. Hence, we propose they represent a response of the protein that correlates with the photoresponse of bound AdoCbl.

### The CarH protein stabilises the productive MLCT state of AdoCbl

As mentioned above, the MLCT state spectrum observed previously in UV-visible TA data from CarH^5^ resembles that of free MeCbl^12^ and AdoCbl bound to its dependent enzyme, glutamate mutase.^26^ Calculations suggest significant mixing between the 2p_z_ orbital of the upper axial ligand and the 3d_z_^2^ of the Co in MeCbl,^27^ which presumably facilitates the formation of a MLCT state where charge density is transferred to the alkyl ligand. Molecular orbital calculations of the MLCT state of photoexcited AdoCbl in glutamate mutase suggest it is dominated by Co d-orbital / corrin π to corrin π* transition.^28^ In both MeCbl^12^ and glutamate mutase-bound AdoCbl,^26^ however, this state decays within ∼ 1 ns and ∼ 100 ps, respectively, to leave the cob(II)alamin / alkyl radical pair. By sharp contrast, in CarH this state is much longer-lived and the UV-visible TA data show no evidence of subsequent radicals.^5, 11^ Interestingly, the EAS from global analysis of TRIR data from MeCbl reveal a sequence of components that are highly similar to those from free AdoCbl (Fig. S13). Hence, the MLCT state doesn’t appear to have a distinctive vibrational signature in the TRIR data following photoexcitation of MeCbl.^14^

In contrast to MeCbl, the TRIR signals in the third EAS from CarH are distinctive (Fig. 2b). This fact, and the outcome of the heating experiments (Figs. S10-12), suggest that the CarH signals in the amide I arise from the protein and not the chromophore. If so, the separation of this protein signal from those from AdoCbl could provide an alternative explanation for why they are convoluted with the radical signals from the cofactor in the second EAS (Figs. 2b&S9a). The kinetics of this protein signal correlate very strongly with those of the MLCT state signal from the UV-visible data^5^: they first appear within 10-15 ps (Fig. 1c) and have a lifetime of 2.94 μs. This lifetime is consistent with the proposal by Miller *et al*. that intersystem crossing in the MLCT populates a six-coordinate triplet state, which would last for enough time to allow for photoproduct formation,^11^ presumably *via* concerted Co−C bond heterolysis and β-hydride elimination. Our TRIR signals represent strong evidence that the CarH protein stabilises the MLCT state and putative triplet state of bound AdoCbl to steer the photochemistry towards this productive channel without releasing reactive radical species.

Computational studies give some indication of how this stabilisation might be achieved. Time-dependent density functional theory suggests the peak of the α/β UV-visible absorption band of the AdoCbl bound to CarH is associated with the S_3_ excited state, which is predominantly MLCT in character.^29^ Molecular dynamics simulations and hybrid quantum mechanical / molecular mechanical calculations propose two interactions between AdoCbl and the protein that could serve to stabilise the MLCT state (Fig. S14).^30^ First, π-stacking between the adenine ring of AdoCbl and Trp131 stabilises CT to the Ado and this stacking appears to be supported through interaction between an adjacent Glu141 to both Ado and Trp131. Second, a H-bond between Glu 175 and His177 that serves as the lower axial base of the cobalamin promotes CT to the upper axial ligand. Variants of CarH containing appropriate mutations at these positions will be investigated as part of future TRIR studies.

In conclusion, our data enable us to chart the course of electron density across AdoCbl following its photoexcitation in CarH (Fig. 3b). This results in photodissociation of the upper axial Ado while avoiding release of radical photoproducts, which CarH achieves by stabilising a series of CT states of AdoCbl. Electron density initially shifts from the corrin to the Co (LMCT), perhaps with some transferred directly from the corrin to the upper axial Ado (LLCT). The charge is then transferred from the Co to the Ado, giving the bond some dative character. The resulting MLCT state is stabilised by the protein for sufficient time (μs) for what is likely to be concerted heterolytic Co−C bond breaking / β-hydride elimination from the Ado. This study provides motivation for a more detailed analysis of the role of the specific residues in stabilising the MLCT state.

**Fig. 3.**
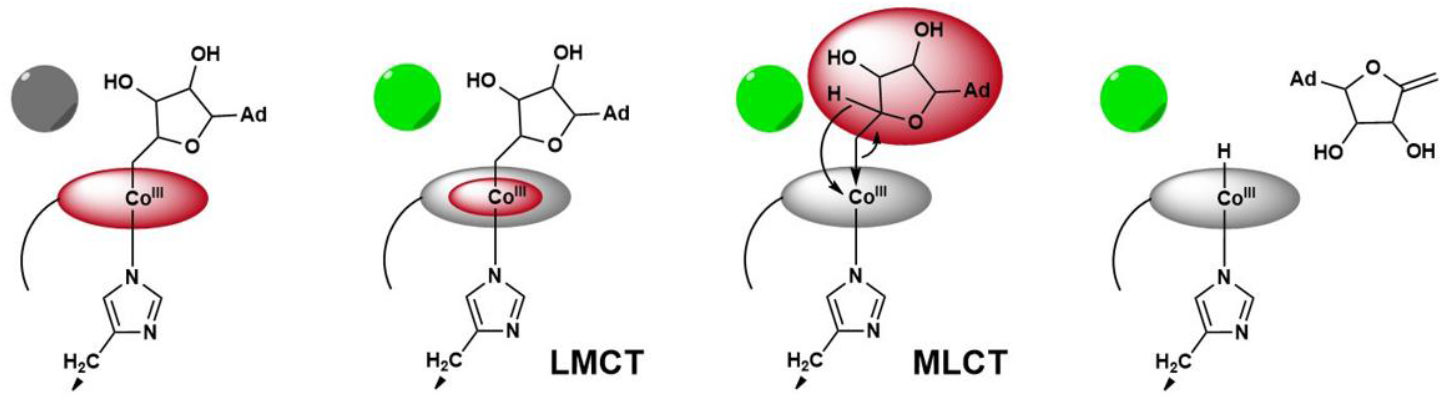
Model showing transfer of electron density (red) to different CT states of AdoCbl following photoexcitation of CarH with light (green circle).

This work was supported by: STFC-funded access to the Ultra Facility; the National Measurement System of the UK Government Department for Science, Innovation, and Technology; a BBSRC CASE studentship to EW (ref 2777745).

## Supporting information

Electronic Supplementary Information

