## Supplementary material for "Tuning of B_12_ photochemistry in the CarH photoreceptor to avoid radical photoproduct": Electronic Supplementary Information

**Supplementary Figures**

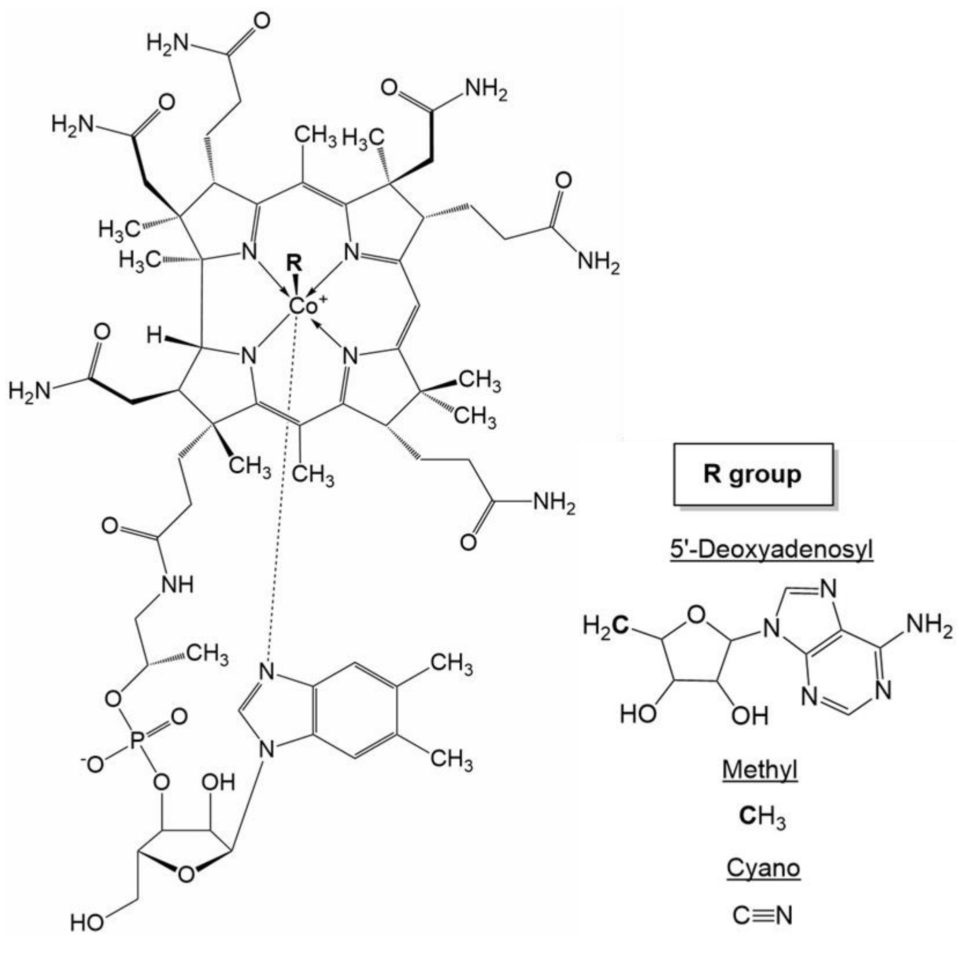


**Fig. S1. Vitamin $B_{12}$ Derivatives**. Vitamin $B_{12}$ is a complex cobalt corrinoid (cobalamin). In the free cofactors, a substituent of the tetrapyrrole corrin macrocycle – 5,6-dimethylbenzimidazole – serves as lower axial ligand. In some $B_{12}$-dependent proteins, including in the CarH holoprotein (holoCarH), this is displaced by a histidine residue from the protein (Fig. S5). The upper axial ligand (R) is variable. In this study, we investigated 5'-deoxyadenosylcobalamin (AdoCbl), methylcobalamin (MeCbl) and cyanocobalamin (CNCbl).

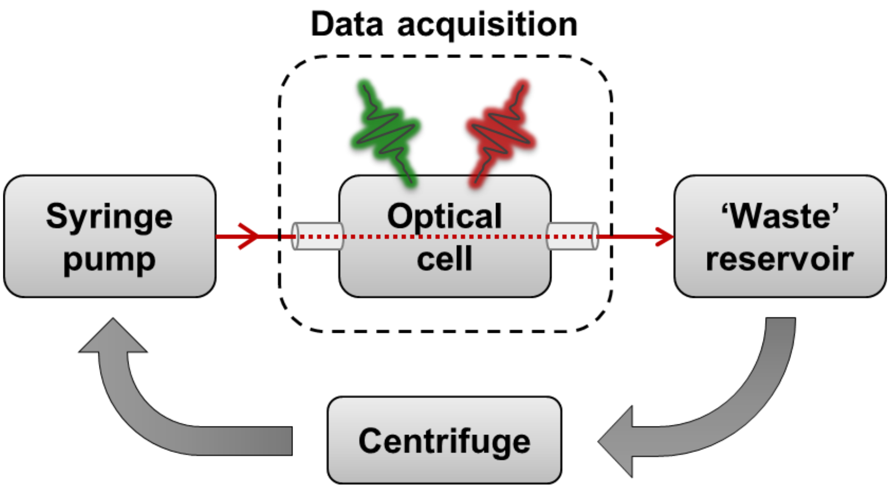


**Fig S2. Sample Flow System for TRIR Measurements.** A bespoke sample flow system was designed and optimised to enable data averaging with holoCarH (*i.e.*, AdoCbl-bound CarH) without the outcome either being confounded by signal from soluble photoproduct or obscured by scatter from precipitated photoproduct. Please refer to the Central Laser Facility Annual Report 2020[1] and the Supplementary Methods below for full details. Briefly, 1-2 mL of the sample was flowed from a syringe pump through the optical cell during data acquisition (UV-visible pump: green; mid-IR probe: red) so that fresh sample was excited with each laser flash. The sample was then collected in a 'waste' reservoir, and any precipitated photoproduct removed *via* centrifugation. The supernatant was reloaded into the syringe and flowed again through the system for further data acquisition.

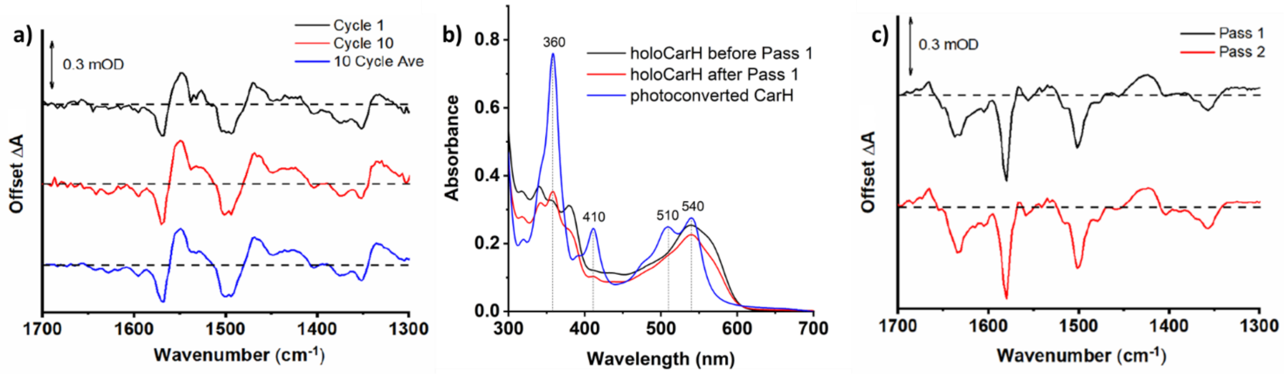


**Fig. S3. Testing of the Flow System. a)** TRIR difference spectra at 2 ps delay following photoexcitation of 360 μM AdoCbl at 525 nm. This concentration of AdoCbl was chosen because it is of a similar magnitude to the concentration of holoCarH used for TRIR. For full dataset, see Fig. S4. Data included are from the first (black) and last (red) data acquisition cycles for a given 'pass' of sample through the flow system in Fig. S2 and the data averaged across all 10 passes (blue). There is minimal signal variation across the 10 cycles. **b)** UV-visible absorption spectra comparing holoCarH before and after the first 'pass' through the flow system with the fully photoconverted spectrum. A small amount of soluble photoproduct (*i.e.*, Cbl-bound CarH) is evident following the first pass, but the spectrum remains dominated by the dark state of CarH (*i.e.,* AdoCbl-bound CarH). **c)** Averaged TRIR 2 ps TRIR difference spectra following photoexcitation of ~250 μM CarH at 525 nm, from both the first (black) and second (red) pass through the flow system. Despite evidence of a small accumulation of soluble photoproduct following the first pass in (b), the TRIR are remarkably consistent between the data sets. This confirms that the signal is dominated by the response of the CarH dark state in each case.

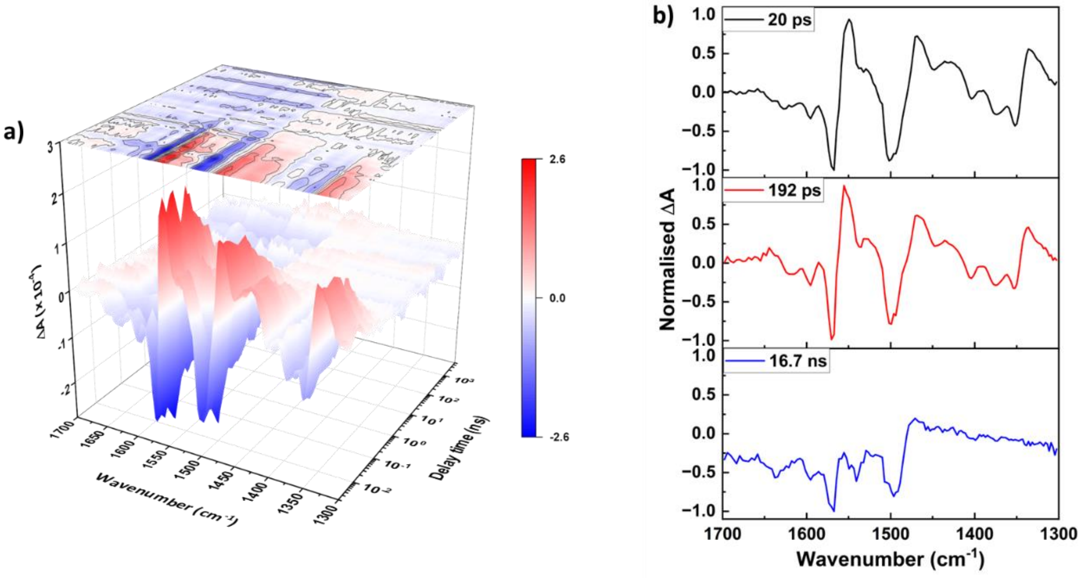


**Fig. S4. TRIR Data from 360 μM AdoCbl at 525 nm**. Free $B_{12}$ derivatives are soluble in aqueous solution up to low mM concentrations, depending on the properties of the upper axial ligand. Most TRIR data for AdoCbl, MeCbl and CNCbl were therefore acquired at 3 mM in buffered $D_2O$ to maximise the S/N and hence the data quality. Because it was only possible to acquire TRIR data from holoCarH at sub-mM concentration, however, we acquired an equivalent dataset over 2 ps – 5 μs following photoexcitation at 525 nm for AdoCbl **(a)** and analysed globally to produce EAS **(b)**. Also see data from the 2 ps delay at this concentration in Fig. S3a. These data confirm consistent, albeit noisier, signals to those observed at higher concentrations (Fig. 2b). The EAS are almost identical and, although their lifetimes are consistently slightly longer, they are of a very similar order.

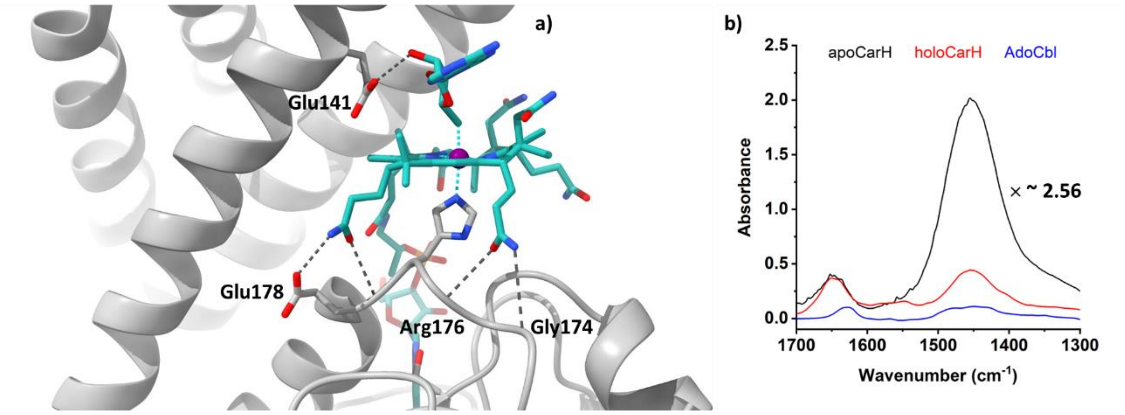


**Fig. S5. AdoCbl Binding to CarH. a)** Inspection of the published crystal structure (PDB:5C8D) of CarH[2] reveals a relatively modest network of H-bonds between the protein and AdoCbl, which is otherwise quite solvent exposed. Only two of the corrin propionamides, both on the same side of the macrocycle, interact directly with CarH: one with both the side chain and backbone of Glu178, and the other with the backbone regions of Arg176 and Gly174. There is a single H-bond between Glu141 and a ribose OH from the upper axial Ado. **b)** For clarity, the FTIR spectra in Fig. 2a are normalised to the highest amplitude peak (*i.e.*, the amide II' band). An unintended consequence of this is that the apoCarH amide I appears of relatively low amplitude. When the apoCarH (90 μM) and holoCarH (230 μM) spectra are normalised to concentration (*i.e.*, the apoCarH amplitude multiplied by ~ 2.56), however, it is clear that the protein amide I signals are of the same relative amplitude. The FTIR spectrum of 3.4 mM AdoCbl is also plotted for comparison. These spectra support our conclusion that the amide I band from holoCarH is dominated by signal from the protein.

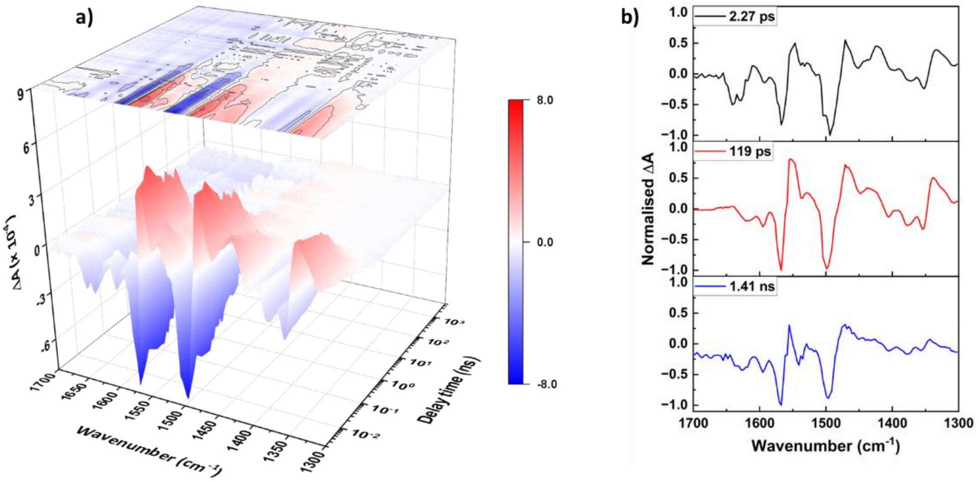


**Fig. S6. TRIR Data from AdoCbl at 380 nm.** To investigate the wavelength dependence of the AdoCbl photoresponse, TRIR data were acquired over 2 ps – 5 μs following photoexcitation at 380 nm of a 3 mM solution of AdoCbl in buffered $D_2O$ **(a)** and analysed globally to produce EAS **(b)**. The data, EAS and their lifetimes are all very similar to those acquired following photoexcitation at 525 nm (Figs. 1b&2a).

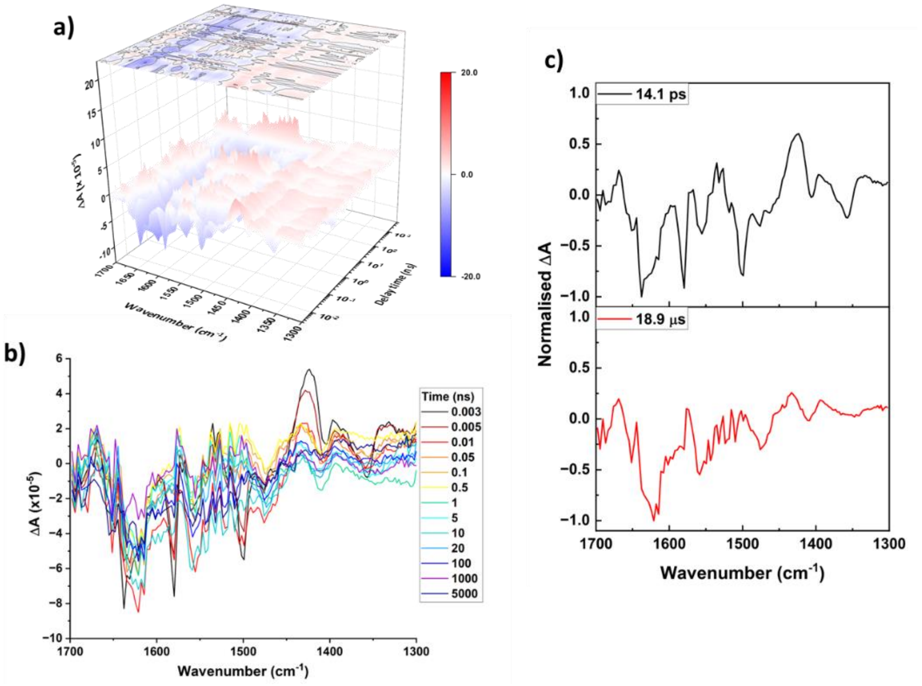


**Fig. S7. TRIR Data from holoCarH at 380 nm.** To investigate the wavelength dependence of the holoCarH photoresponse, TRIR data were acquired over 2 ps – 5 μs following photoexcitation at 380 nm of a ~250 μM solution of holoCarH in buffered $D_2O$ **(a&b)** and analysed globally to produce EAS **(c)**. Owing to a large signal artefact at the 2 ps delay time, for clarity we only plot data in (a&b) from 3 ps. The data reveal a significantly reduced amplitude relative to the equivalent data acquired following photoexcitation at 525 nm (Fig. 1c). This means the features are difficult to compare when plotted in 3D (a), but some are apparent when plotted in 2D (b), not least the long-lived bleach and transient in the amide I and II' regions, respectively. Global analysis only produces two EAS above the noise (c), in contrast to the three from the 525 nm data (Fig. 2b). Despite the noise, these two components and their lifetimes resemble closely the second and third EAS from the 525 nm data. Because of the poor S/N evident in (a&b) and the signal artefact at early time, it was not possible to resolve the first component. We conclude that the main difference between photoexcitation of holoCarH in the green and UVA is the productive quantum yield, consistent with reports elsewhere.[3]

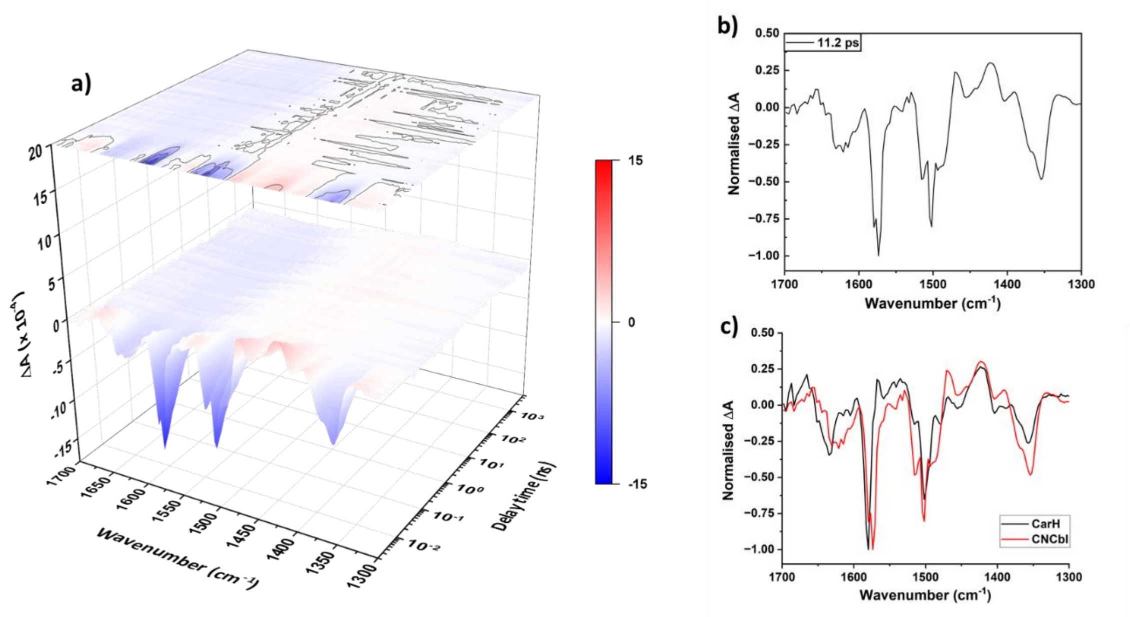


**Fig. S8. TRIR Data from CNCbl at 525 nm.** TRIR data were acquired over 2 ps – 5 μs following photoexcitation at 525 nm of a 3 mM solution of CNCbl in buffered $D_2O$ **(a)** and analysed globally to produce EAS **(b)**. The data are consistent with those we reported previously,[4] and again produce a single EAS as expected. This component represents an excited state with predominantly a ligand-to-metal (corrin to Co) charge transfer (LMCT) character.[5, 6] When overlaid **(c)**, it is clear that it shares many of the features of the first EAS from the global analysis of the holoCarH TRIR data (Fig. 2b), with only minor variations in the frequency and relative amplitude of the peaks.

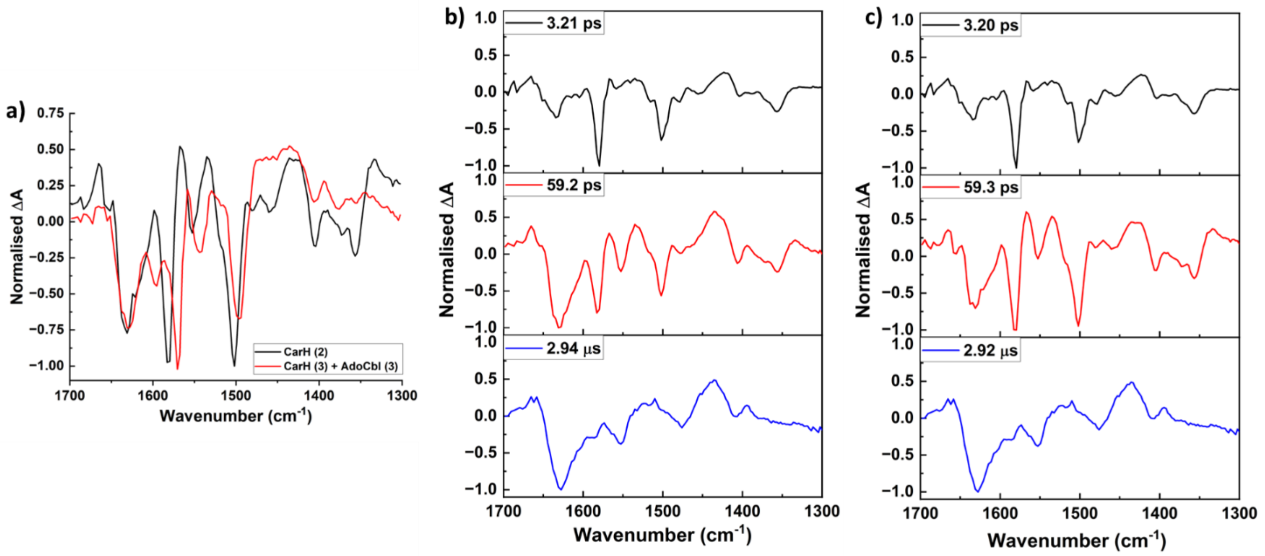


**Fig. S9. Signal Convolution and Analysis Model.** The second EAS from global analysis of holoCarH data following photoexcitation at 525 nm (Fig. 2b) appears to be the convolution of signals from two species: the cob(II)alamin radical (second EAS from free AdoCbl, Fig. 2a) and the longer-lived species represented by the third EAS from holoCarH (Fig 2b). To investigate, we combined these EAS in **(a)**. The combination (red) that produced a spectrum most similar to the second EAS from the holoCarH data (black) was 29:71, respectively, which compares well with the branching ratios (21% radical and 79% productive channel) estimated by Kutta and coworkers.[7] Two global analysis models were therefore considered for the holoCarH TRIR data, one assuming a sequential mechanism **(b)**, the other a branched mechanism **(c)**. Both returned very similar EAS and lifetimes. See main text for discussion and below for sensitivity analysis.

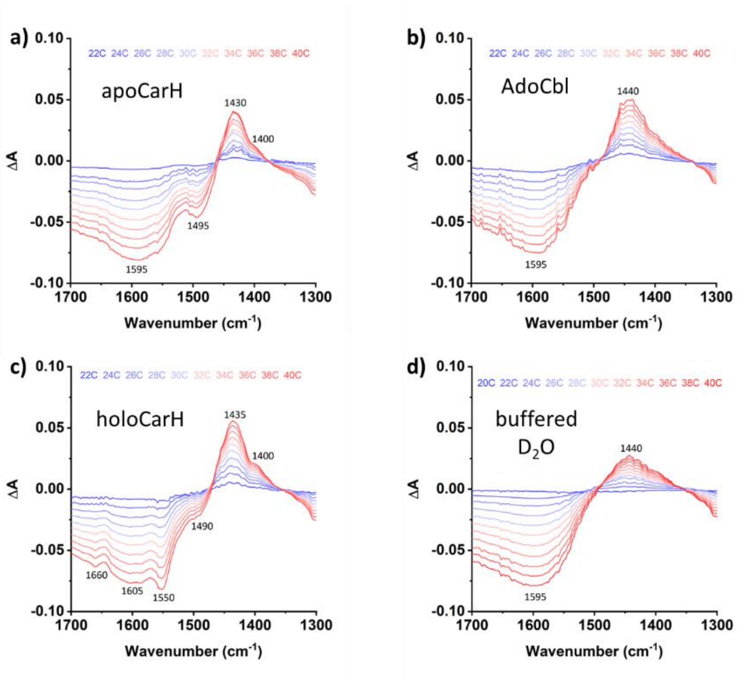


**Fig. S10. Variable Temperature FTIR Spectroscopy 1: Difference Spectra.** To investigate the effect on the IR signals of temperature dissipation through AdoCbl and CarH, FTIR spectra were acquired in buffered $D_2O$ between 1700 – 1300 $cm^{-1}$ for every 2 degrees between 22-40 °C. A corresponding spectrum acquired at 20 °C was then subtracted from the spectrum at each temperature to produce difference spectra for 90 μM apoCarH **(a)**, 300 μM AdoCbl **(b)**, and 230 μM holoCarH **(c)**. To identify the signal changes from the solvent, equivalent data were acquired from $D_2O$ **(d)**. These were used for further data processing (Fig. S11).

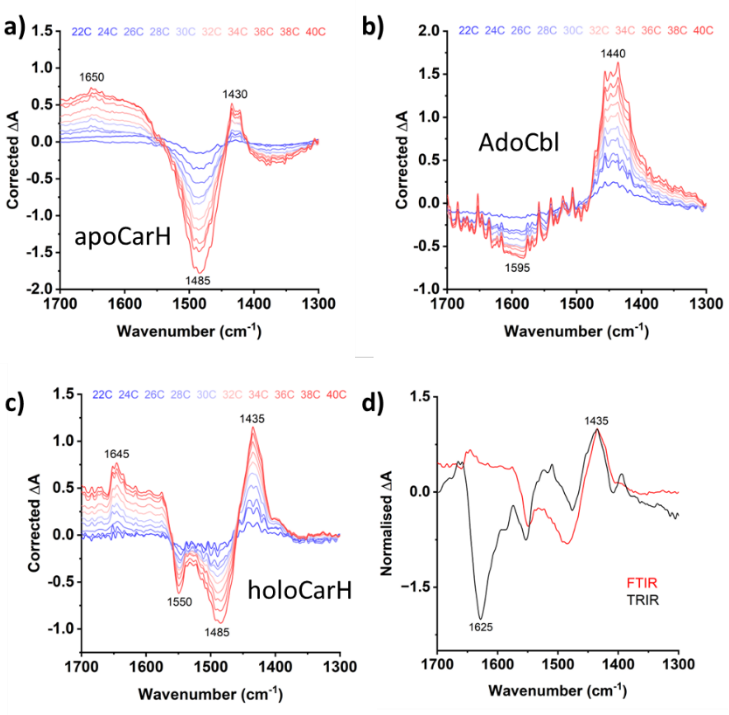


**Fig. S11. Variable Temperature FTIR Spectroscopy 2: Processed Difference Spectra.** The difference spectra in Fig. S10 were processed first by normalising the largest amplitude temperature-induced change (which is from the 40 - 20 °C difference spectrum) to 1 and scaling the difference spectra calculated at all other temperatures accordingly. The normalised difference spectrum of $D_2O$ at each temperature was then subtracted from the equivalent data for each sample to produce processed difference spectra that reflect the temperature-induced changes in apoCarH **(a)**, AdoCbl **(b)**, and holoCarH **(c)** corrected for solvent effects. The largest amplitude temperature-induced difference spectrum from the holoCarH data was then normalised to a common peak (the positive peak at 1435 $cm^{-1}$) in the third EAS from global analysis of the holoCarH TRIR data (Fig. 2b), and the spectra overlaid **(d)**. This overlay helps identify which peaks in the third EAS might have contributions from heat dissipation (see main article for discussion).

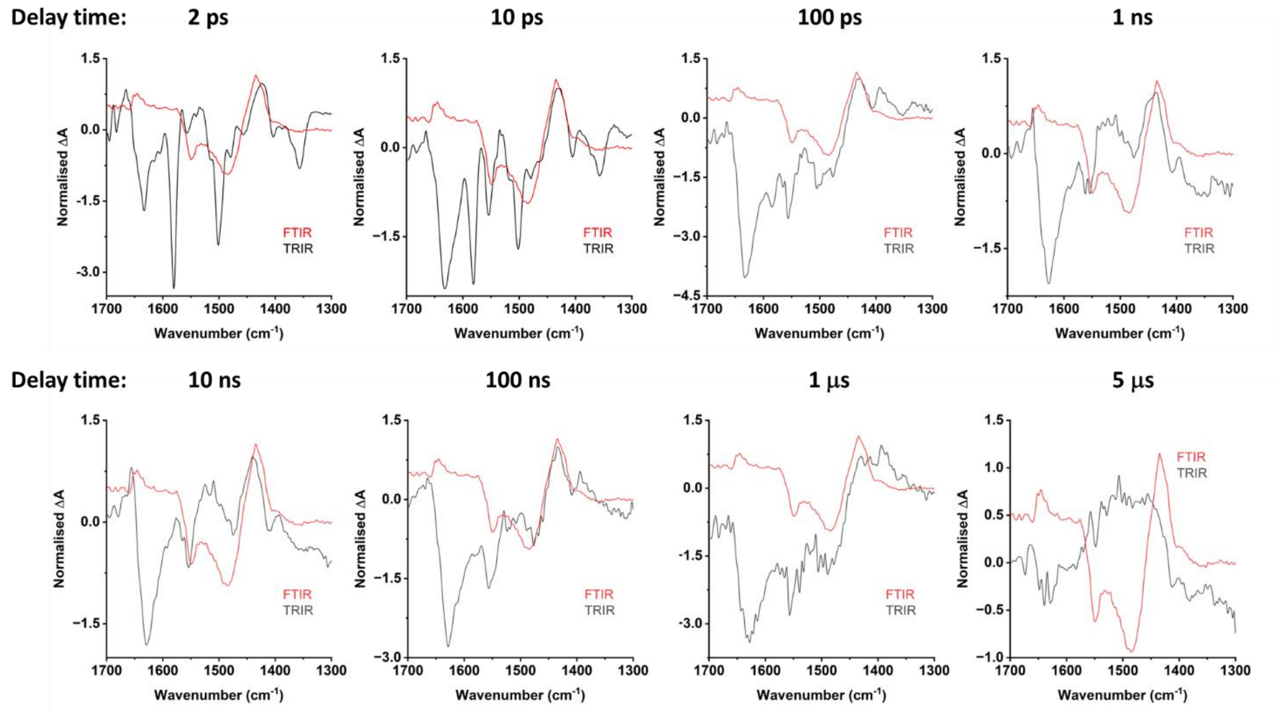


**Fig. S12. Variable Temperature FTIR Spectroscopy 3: Comparison as a Function of Time.** To further compare the temperature induced changes in the CarH FTIR signals with the TRIR data, the FTIR difference spectra were normalised as described in Fig. S11d and overlaid with TRIR difference spectra from the holoCarH data at eight selected delay times between 2 ps and 5 μs. Spectral features in the TRIR spectra that are similar to those observed in the variable temperature difference spectrum (the transient at ~ 1430 $cm^{-1}$ and bleaches at ~ 1490 $cm^{-1}$ and 1555 $cm^{-1}$) appear to decay over a ~1 μs timescale. The bleach at ~ 1640 $cm^{-1}$ in the amide I region cannot be accounted for, however, and is therefore assigned to protein signal relevant to the productive CarH photoresponse. At 5 μs, the absolute amplitude of the TRIR data becomes very low (it appears larger here because of the normalisation), but some vestiges of the bleach from the protein at ~ 1640 $cm^{-1}$ remain.

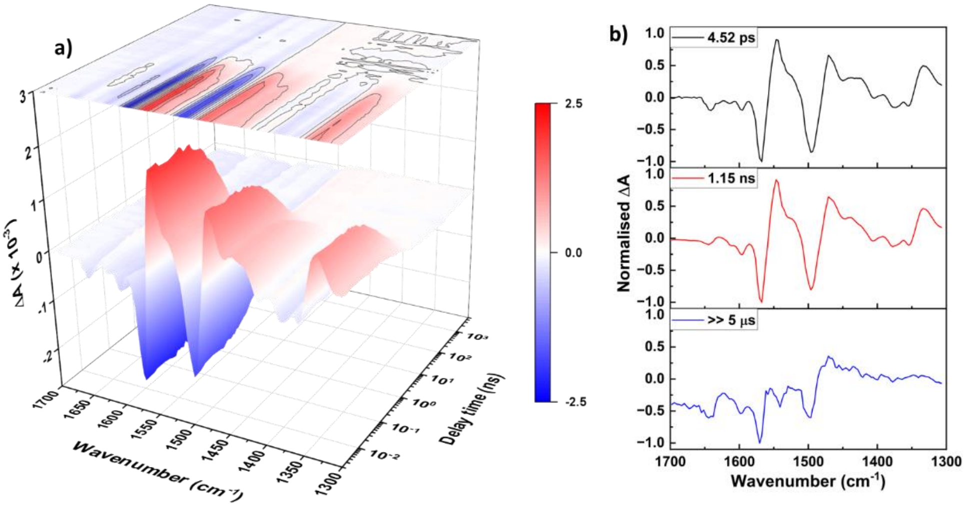


**Fig. S13. TRIR Data from MeCbl at 525 nm.** TRIR data were acquired over 2 ps – 5 μs following photoexcitation at 525 nm of a 3 mM solution of MeCbl in buffered $D_2O$ **(a)** and analysed globally to produce EAS **(b)**. The data are consistent with those we reported previously.[4] Although photoexcitation of MeCbl is known to produce a metal-to-ligand charge transfer (MLCT) state before dissociating into radicals, this state does not appear to have a distinctive vibrational signature. These data and EAS instead closely resemble those from AdoCbl, albeit with different kinetics. Global analysis of these data returned a lifetime for the 3rd component (bottom panel) of ~ 36 μs; however, because our data acquisition window is 5 μs, the uncertainty associated with this is high.

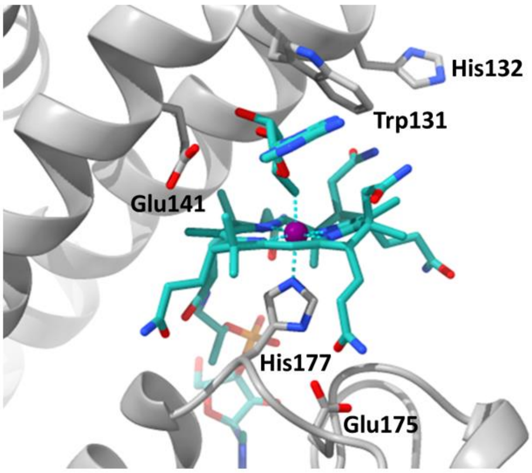


**Fig. S14. Residues in CarH proposed to stabilise the MLCT state.** Various residues around the upper and lower axial ligand of AdoCbl have been proposed by Cooper *et al*.[8] to stabilise the MLCT state. See main article for discussion. Some of these will be targeted for mutagenesis for future TRIR studies.

**Supplementary Methods**

### Materials

All commercial reagents – coenzyme $B_{12}$ (5'-deoxyadenosylcobalamin, AdoCbl), methylcobalamin (MeCbl), vitamin $B_{12}$ (cyanocobalamin, CNCbl), $D_2O$ – were obtained from *Sigma-Aldrich*, of analytical grade, and used without further purification.

#### Protein overexpression and purification

The plasmid containing the gene encoding wild-type CarH from *Thermus thermophilus* was kindly provided by S. Padmanabhan and Montserrat Elías-Arnanz. The gene had been cloned into a modified pET15b (*Novagen*) expression vector using NdeI and BamHI restriction enzymes, to provide a N-terminal 6xHis affinity tag as previously described.[9]

The plasmid was transformed into *E. coli* BL21 (DE3) cells (*Novagen*), and a single colony inoculated into a small volume of selective ($Amp^R$) LB medium. The cell cultures were grown to an $OD_{600}$ 0.8-1.0 at 37 °C and 200 rpm, at which time they were inoculated into a larger volume of fresh LB. The fresh cell cultures were grown to an $OD_{600}$ 0.6-0.8 at 37 °C and 200 rpm, at which time they were cooled down to the induction temperature of 25 °C. Protein expression was then induced with 0.5 mM IPTG and the cells left to incubate overnight at 25 °C and 200 rpm. Cells were then harvested by centrifugation at 6,000 x g and 4 °C for 20 mins, collected, rapidly frozen in liquid nitrogen, and stored at - 80 °C. For purification, cells were thawed and resuspended in buffer A (50 mM sodium phosphate + 300 mM NaCl, pH 7.5) supplemented with 2.5 mM imidazole, lysozyme, DNAse, $MgCl_2$, and protease inhibitors, and lysed by cell disruption (30 kpsi, 2 passes). Cell debris was removed by centrifugation at 48,000 x g and 4 °C for 1 h. The collected supernatant was filtered through a 0.22 µm membrane and incubated with 5 mL TALON metal affinity resin (*Clontech*) for 2 h at 4 °C with rolling. Protein-bound resin was washed with buffer A (see above) supplemented with 5 mM (wash 1) and 10 mM imidazole (wash 2). The CarH apoprotein (apoCarH) was eluted with buffer A supplemented with 150 mM imidazole following 1 h of incubation at 4 °C with rolling. Fractions containing purified protein were further purified by size-exclusion chromatography using a Superdex200 high-performance liquid chromatography column (*Cytiva*), equilibrated with 50 mM sodium phosphate buffer + 150 mM NaCl, pH 7.5. Purified apoCarH was brought to the desired concentration using 10 kDa molecular weight cut off Vivaspin centrifugal filter devices (*Sartorius*).

#### Protein preparation for spectral measurements

Because $H_2O$ absorbs strongly in the IR region of interest (1700 – 1300 $cm^{-1}$), all samples were prepared in buffered $D_2O$ (50 mM sodium phosphate buffer + 150 mM NaCl, adjusted to pH 7.91 (pD 7.5) with NaOD).

A solution of purified apoCarH (< 150 µM to remain soluble) in buffered $D_2O$ was incubated with a 5-fold excess of AdoCbl in buffered $D_2O$ for 15 mins at 4 °C. Unbound AdoCbl was then removed using a size-exclusion column (*Bio-Rad*) pre-equilibrated with buffered $D_2O$ (as above), which resulted in a 1.5-fold dilution. The holoCarH was brought up to working concentration (200-300 µM) using centrifugal filter devices. The entire process was conducted under red light (> 600 nm).

#### Static spectroscopy

UV-visible absorption spectra of AdoCbl, MeCbl, CNCbl, apoCarH and holoCarH were acquired between 250 – 700 nm, using a Cary 60 spectrophotometer (*Agilent Technologies*) and a 1 mm quartz cuvette. Background spectrum was collected with buffered $D_2O$.

FTIR absorption spectra of AdoCbl, MeCbl, CNCbl, apoCarH, and holoCarH were acquired using a Nicolet iS20 FTIR spectrometer (*Thermo Scientific*). Although $H_2O$ contamination signals can be mitigated significantly by using $D_2O$, very short pathlengths must still be used in case of any residual $H_2O$. Thus, a sample cell with $CaF_2$ windows and a PTFE spacer to give a pathlength of 100 µm pathlength was used (see 'TRIR spectroscopy' section for a detailed description of the cell), which allowed acquisition of signals in the mid-IR region (4000 – 400 $cm^{-1}$). A background spectrum was collected with buffered $D_2O$, and 100 scans/measurement were acquired for all samples. Measurements with photosensitive samples were conducted under red light (> 600 nm).

To test the heat dissipation hypothesis, spectra were acquired between 1700 – 1300 $cm^{-1}$ every 2 degrees between 20-40 °C, using a thermostatic Harrick cell with $CaF_2$ windows and a 100 µm pathlength, connected to a water bath and a temperature controller unit (*Harrick Scientific*). All samples were equilibrated for 2 minutes after the temperature set with the water bath had stabilised on the temperature control unit display.

**TRIR spectroscopy**

TRIR spectroscopy was performed using the Ultra A setup in TRMPS mode at the Central Laser Facility, STFC, Rutherford Appleton Laboratory, UK. This uses a 100 kHz ultrafast laser based on a custom dual Yb:KGW system (*Pharos, Light Conversion*). Datasets were acquired for AdoCbl, MeCbl, CNCbl, and holoCarH following photoexcitation at 525 nm (corresponding to the $\alpha\beta$ absorption band). Data were additionally acquired for AdoCbl and holoCarH following photoexcitation at 380 nm ($\gamma$ absorption band) to investigate any dependence of the photoresponse on excitation wavelength. An estimate of the absorbance in the green at the working concentrations of holoCarH (200-300 µM) and free $B_{12}$ derivatives (3 mM) in a 100 µm pathlength are ~ 0.02-0.03 and 0.3, respectively. The optical absorption by the samples is therefore linear and avoids inner-filter effects under these conditions. The excitation pump laser was set with a pulse energy of 1 µJ at 1 kHz repetition rate, and the polarization of it was set at 54.7° with respect to the IR probe beam. All data were acquired under red light (> 600 nm) to stop spurious photoactivation from the ambient lighting.

The measurements were conducted at room temperature using a purpose-built sample flow system,[1] where each sample (~1-2 mL) was uni-directionally flowed using a syringe pump through a modified small Harrick cell (with 2 mm thick $CaF_2$ windows and a 100 µm spacer) to 'waste' (Fig. S2). The dead volume required for the sample to reach the optical window (tubing length / diameter, cell attachments, window area, etc.) was minimized while still enabling bubble-free, uniform flow and for the flow setup to interface with the optical setup while accounting for rastering (*i.e.*, moving) of the cell in the plane perpendicular to the beam. The cell was rastered during data acquisition using a Lissajous pattern to both limit photodegradation and further improve coverage by the laser of the flowing sample. An optimised flow rate of 0.9 mL $h^{-1}$ was found to maximise coverage by the laser of the sample when within the optical window, whilst enabling fresh sample to be excited by each laser pulse and limiting photodegradation between each scan. This setup enabled data acquisition over 10 cycles for ~ 45 minutes (with an averaging time of 7.5 s per delay time) without any significant signal deterioration between cycles owing to photoproduct precipitation. It was also possible to recycle and reuse the samples by collecting the flow-through ('waste') and removing the precipitate (if any) by centrifugation. Indeed, this method successfully removed photoconverted sample and retained a sufficient volume and concentration of holoCarH (confirmed by acquiring static UV-visible spectra, Fig. S3) to acquire a second data set using the sample parameters described above (Fig. S3). This means data from two 10-scan passes could be averaged to improve signal-to-noise ratio. Difference spectra were generated relative to the ground state in the spectral window 1700 – 1300 $cm^{-1}$ at delay times ranging from 2 ps to 5 µs. Pixel to wavenumber calibration was performed using a polystyrene standard.

**Data analysis**

Global analysis was performed using Glotaran software (version 1.5.1) to extract evolution associated spectra (EAS) and associated lifetimes.[10] Glotaran enables a singular value decomposition (SVD) approach to modelling the data and resolves the 3D matrix encompassing absorbance, wavelength and time into a series of kinetic components which decay exponentially. The EAS are plots of the pre-exponential amplitude for each component against wavelength, representing the different transient species resolved in analysis.

Datasets were fitted using both sequential and branched analysis schemes (Fig. S9). The number of kinetic components was chosen to minimise the residual error after fitting (*i.e.*, the root mean squared, RMS) and to ensure that components had corresponding singular vectors with time and spectral elements significantly different from the noise.

**Uncertainty / sensitivity analysis**

The time constants obtained through global analysis have not been quoted with experimental errors. Due to the time-consuming nature of data acquisition and limited instrument time at the CLF Ultra facility, for CarH it was impractical to acquire replicate datasets of the requisite quality to enable global analysis of all the kinetic components of interest and then calculate the standard deviation of the resulting time constants. Glotaran software provides an overall RMS error for the global fitting, but this contains limited information about overall uncertainty. A full uncertainty analysis was therefore not possible.

One can, however, conduct a sensitivity analysis into how uncertainty in the output of a model can be apportioned to different sources of uncertainty in the model input.[11] In our case, the model inputs are the time constants used to fit the experimental data, the output is the fit to the data, and the uncertainty of that output is measured by the RMS. Glotaran cannot run if all inputs are fixed, but one input can be varied systematically while letting the software calculate the others. The following figures (S15 – S20) show the relationship between the varied kinetic components for each system studied and the RMS of the associated fittings. Alongside these, we plot the correlations between a given kinetic component with the other unfixed input components in the model.

The values of the kinetic components extracted by Glotaran when the software is left to run freely correspond to minima in the RMS (*i.e.*, where the fit uncertainty is lowest). To be confident in the kinetic components extracted from global analysis, there should be a steep singular minimum, or multiple minima where one is deeper than the rest. In our analyses, the RMS values depended on whether the model was left to run freely (as with the original fitting), or one parameter was fixed (as in the sensitivity analysis). The latter produced slightly smaller RMS, which may be explained by a reduction in the degrees of freedom. For the analyses to remain physically meaningful, if two components were close in magnitude, we avoided varying the parameter values such that their values crossed over (*e.g.,* if varying $k_1$ and initially $k_1 > k_2$, we would not vary it to the point $k_1 < k_2$).

The RMS plots showed a single minimum for each kinetic component from all data for CarH (Figs. S15&16) and free AdoCbl (Figs. S17-19). Much like the global analysis (Fig. S9), the sensitivity analysis for the branched CarH model gave the same results as the linear sequential model for both the RMS and values of the kinetic components (Fig. S15).

The sensitivity analysis of the MeCbl fitting revealed two minima for each component (Fig. S20). The minimum a given run would converge upon for MeCbl depended on the guesses used as starting values for the input parameters. The time constants corresponding to the deepest RMS minima were therefore presented in this work ($k_1$ = 2.14, $k_2$ = 0.566, $k_3$ = 1.88 $\times$ $10^{-5}$, with units of $ns^{-1}$). However, those associated with both minima for each component, including the values of $k_1$ = 222, $k_2$ = 0.871, $k_3$ = 2.79 $\times$ $10^{-5}$ $ns^{-1}$ corresponding to the shallower minima, agree with the kinetic parameters extracted from other time-resolved studies to a similar extent. For example, Shiang and coworkers obtained a kinetic component of 60 $ns^{-1}$ from UV-visible transient absorption spectroscopy, which lies in between the values of $k_1$ associated with the minima in this work.[11] They also obtained a subsequent component of 1.1 $ns^{-1}$, which resembles both our $k_2$ values at RMS minima, but they did not obtain a value for $k_3$ due to the shorter timescale of their experiment.

A sensitivity analysis was not performed for the CNCbl data because only one component was extracted from the fit and Glotaran cannot run if every input component is fixed. However, different starting values for this input parameter were trialled over a few orders of magnitude (1-1000 $ns^{-1}$). These resulted in the same fit and kinetic components, making multiple minima unlikely.

Because Glotaran could not perform global lifetime analysis if every component was fixed, the kinetic parameters used as inputs for the model were correlated. In general terms, we found that the kinetic input parameters for a given fit were more closely correlated if they were closer in magnitude to one another. The nature of the correlations varied between the different datasets, but a full analysis of this (including correlations with component amplitude) are beyond the scope of this study.

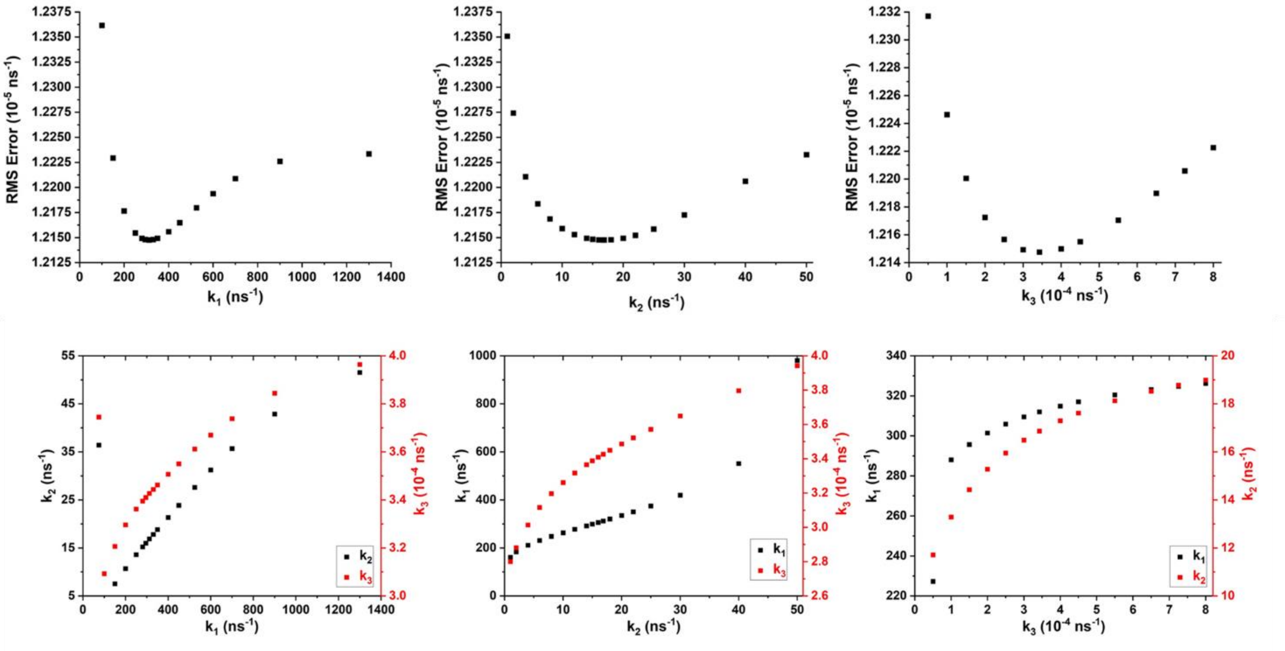


**Fig. S15. Sensitivity Analysis 1: HoloCarH ($\lambda_{ex}$ = 525 nm) – Linear or Branched (Results were Identical).** To investigate the sensitivity of the global analysis fitting of the holoCarH TRIR data ($\lambda_{ex}$ = 525 nm), the associated RMS was plotted as a function of each kinetic component (top), where each component was fixed and then systematically varied. The correlations between the different kinetic components – *i.e.*, the effect of varying one on the values of the others – are also presented (bottom).

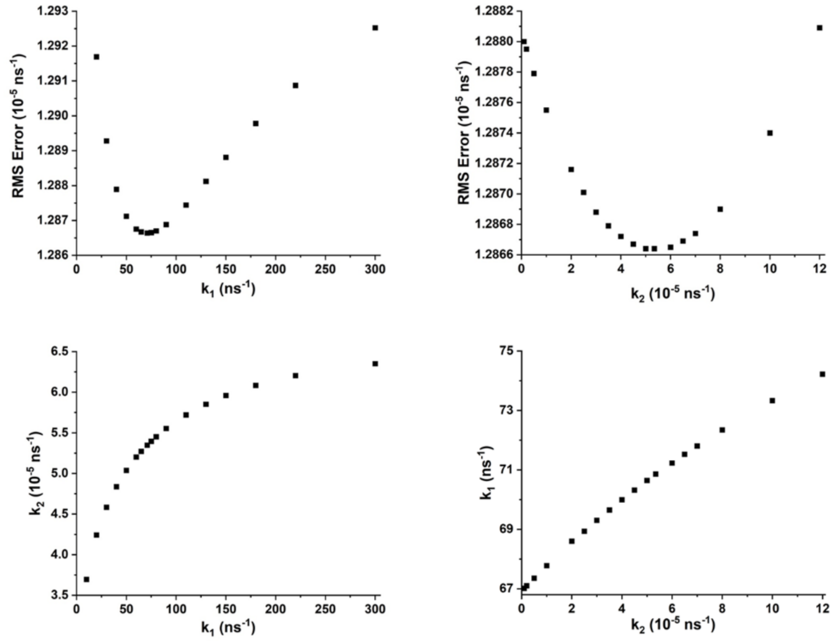


**Fig. S16. Sensitivity Analysis 2: HoloCarH ($\lambda_{ex}$ = 380 nm).** To investigate the sensitivity of the global analysis fitting of the holoCarH TRIR data ($\lambda_{ex}$ = 380 nm), the associated RMS was plotted as a function of each kinetic component (top), where each component was fixed and then systematically varied. The correlations between the different kinetic components – *i.e.*, the effect of varying one on the values of the others – are also presented (bottom).

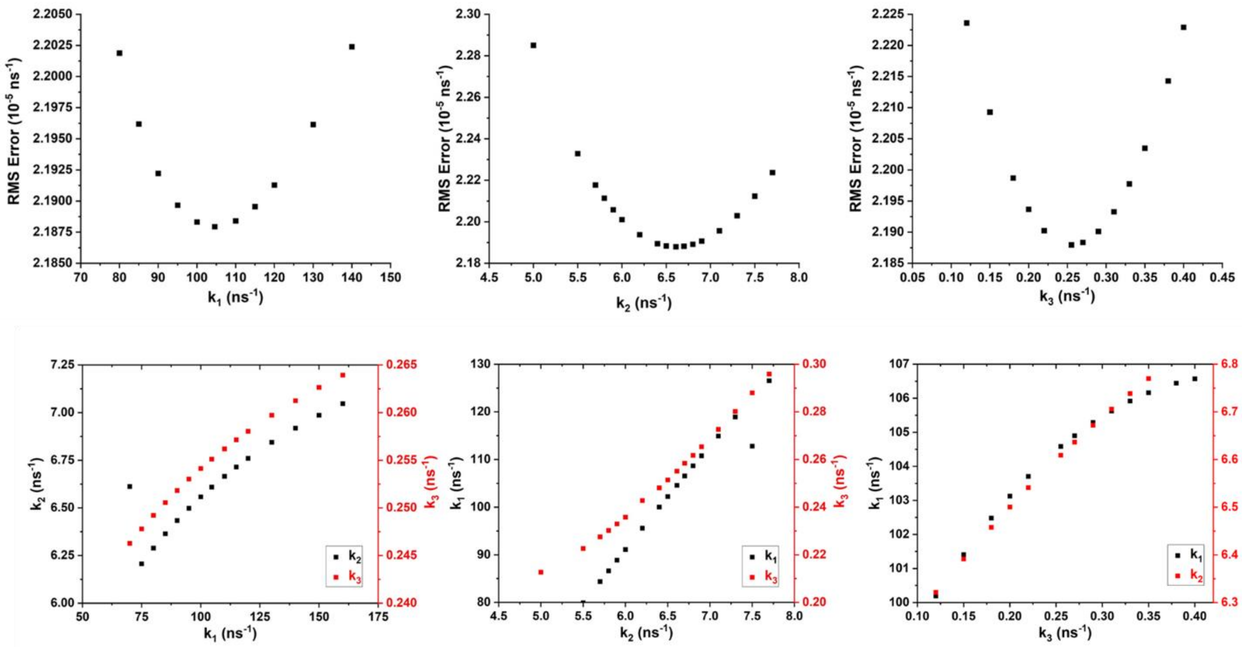


**Fig. S17. Sensitivity Analysis 3: 3 mM AdoCbl ($\lambda_{ex}$ = 525 nm).** To investigate the sensitivity of the global analysis fitting of the 3 mM AdoCbl data ($\lambda_{ex}$ = 525 nm), the associated RMS was plotted as a function of each kinetic component (top), where each component was fixed and then systematically varied. The correlations between the different kinetic components – *i.e.*, the effect of varying one on the values of the others – are also presented (bottom).

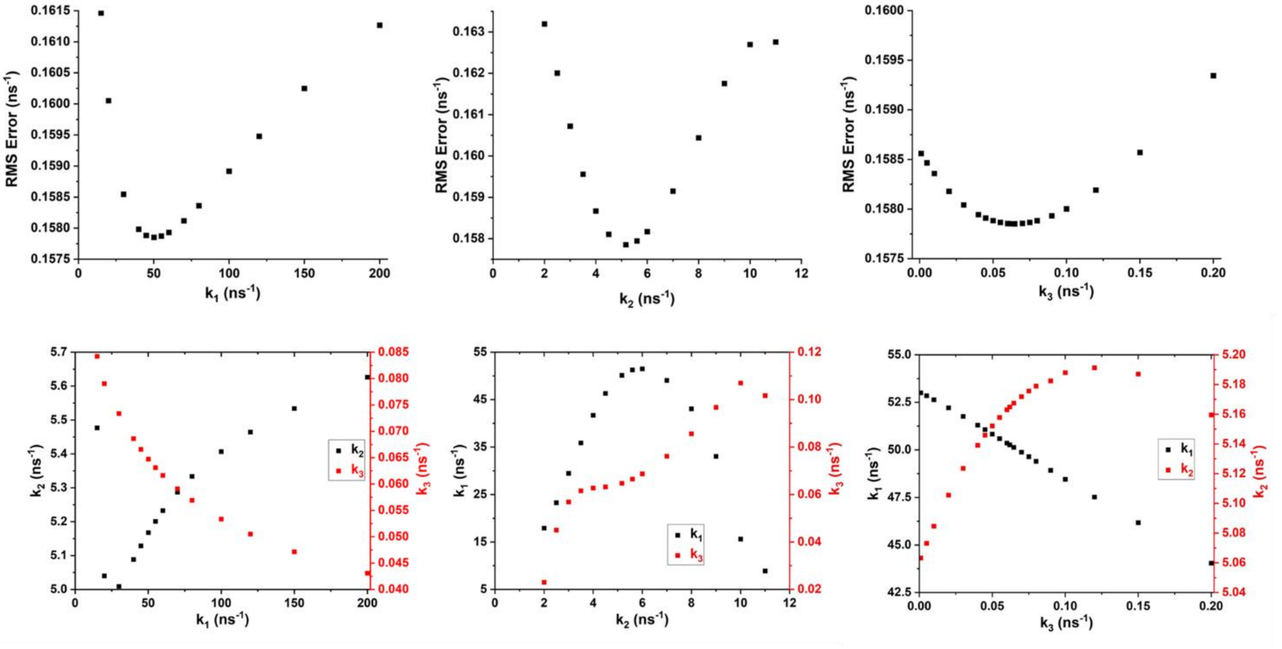


**Fig. S18. Sensitivity Analysis 4: 260 µM AdoCbl ($\lambda_{ex}$ = 525 nm).** To investigate the sensitivity of the global analysis fitting of the 260 µM AdoCbl data ($\lambda_{ex}$ = 525 nm), the associated RMS was plotted as a function of each kinetic component (top), where each component was fixed and then systematically varied. The correlations between the different kinetic components – *i.e.*, the effect of varying one on the values of the others – are also presented (bottom).

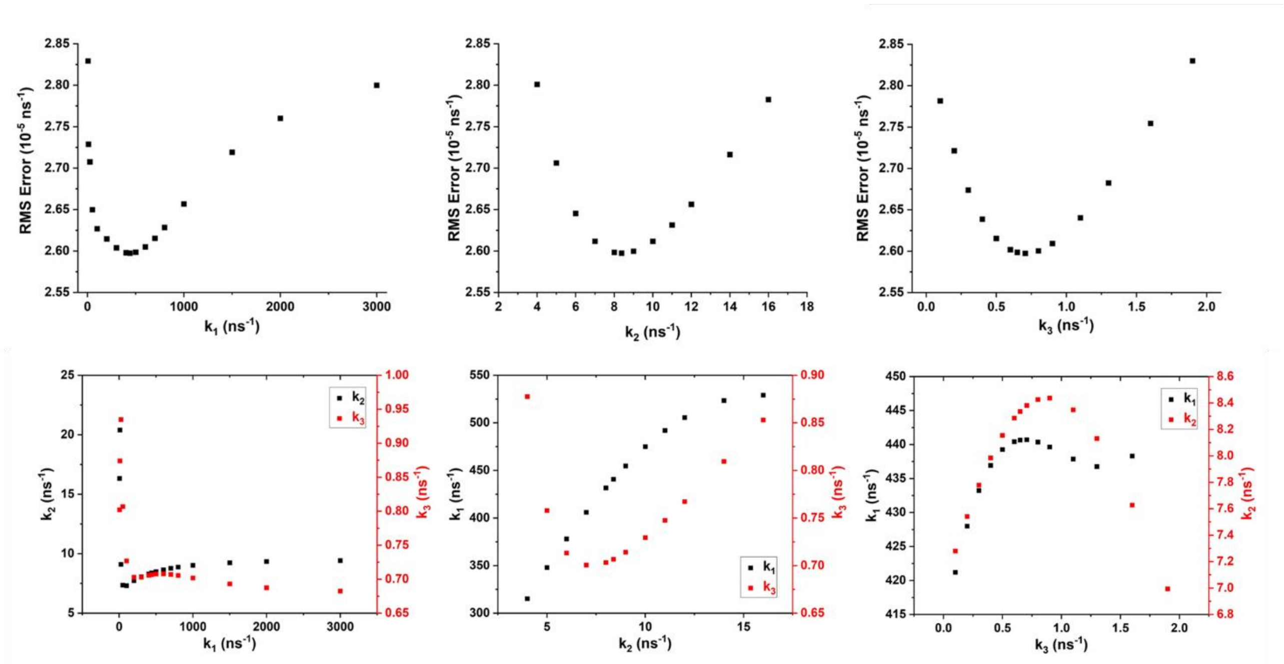


**Fig. S19. Sensitivity Analysis 5: 3 mM AdoCbl ($\lambda_{ex}$ = 380 nm).** To investigate the sensitivity of the global analysis fitting of the 3 mM AdoCbl data ($\lambda_{ex}$ = 380 nm), the associated RMS was plotted as a function of each kinetic component (top), where each component was fixed and then systematically varied. The correlations between the different kinetic components – *i.e.*, the effect of varying one on the values of the others – are also presented (bottom).

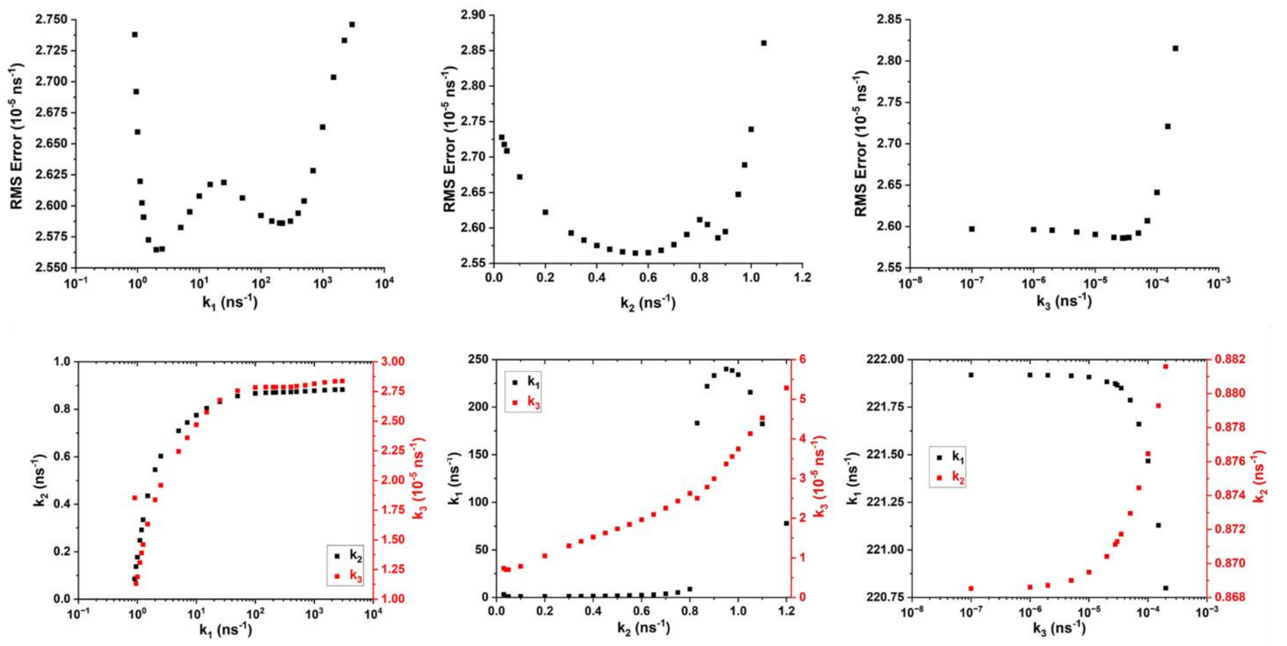

**Fig. S20. Sensitivity Analysis 6: MeCbl ($\lambda_{ex}$ = 525 nm).** To investigate the sensitivity of the global analysis fitting of the MeCbl data ($\lambda_{ex}$ = 525 nm), the associated RMS was plotted as a function of each kinetic component (top), where each component was fixed and then systematically varied. The correlations between the different kinetic components – *i.e.*, the effect of varying one on the values of the others – are also presented (bottom).
